# Sex, gender diversity, and brain structure in children ages 9 to 11 years old

**DOI:** 10.1101/2023.07.28.551036

**Authors:** Carinna Torgerson, Hedyeh Ahmadi, Jeiran Choupan, Chun Chieh Fan, John R. Blosnich, Megan M. Herting

**Author notes:** Corresponding Author: Megan M. Herting, PhD 1845 N Soto Street, Rm 225N Los Angeles, CA 90089. ***Abbreviations***: ABCD, Adolescent Brain Cognitive Development Study; MRI, magnetic resonance imaging; DTI, diffusion tensor imaging; FA, fractional anisotropy; MD, mean diffusivity; ABCD, Adolescent Brain Cognitive Development; WM, white matter; GM, gray matter; T1w, T1-weighted; T2w, T2-weighted; ROI, region of interest; PDS, pubertal development scale.

## Abstract

There remains little consensus about the relationship between sex and brain structure, particularly in childhood. Moreover, few pediatric neuroimaging studies have analyzed both sex and gender as variables of interest - many of which included small sample sizes and relied on binary definitions of gender. The current study examined gender diversity with a continuous felt-gender score and categorized sex based on X and Y allele frequency in a large sample of children ages 9-11 years-old (N=7693). Then, a statistical model-building approach was employed to determine whether gender diversity and sex independently or jointly relate to brain morphology, including subcortical volume, cortical thickness, gyrification, and white matter microstructure. The model with sex, but not gender diversity, was the best-fitting model in 75% of gray matter regions and 79% of white matter regions examined. The addition of gender to the sex model explained significantly more variance than sex alone with regard to bilateral cerebellum volume, left precentral cortical thickness, as well as gyrification in the right superior frontal gyrus, right parahippocampal gyrus, and several regions in the left parietal lobe. For mean diffusivity in the left uncinate fasciculus, the model with sex, gender, and their interaction captured the most variance. Nonetheless, the magnitude of variance accounted for by sex was small in all cases and felt-gender score was not a significant predictor on its own for any white or gray matter regions examined. Overall, these findings demonstrate that at ages 9-11 years-old, sex accounts for a small proportion of variance in brain structure, while gender diversity is not directly associated with neurostructural diversity.

**Highlights:**

- Sex-related variance in regional human brain structure is widespread at ages 9-11
- Together, sex and gender diversity accounted for more variance in only a few regions
- Felt-gender diversity itself was not significantly related to any outcome at ages 9-11
- Effect sizes for sex and felt-gender estimates were small

## 1. Introduction

Although scientific institutions increasingly advise that sex and gender should be disaggregated and studied separately in all human research (Heidari et al., 2016; Mauvais-Jarvis et al., 2020; National Academies of Sciences, 2022; Nielsen et al., 2018; Tannenbaum et al., 2019), few neuroscientific studies to date have included both as variables of interest. As a result, in contemporary human neuroscience literature it remains impossible to parse the extent to which differences between males and females are ascribable to sex versus social-environmental factors like gender (Eliot et al., 2021). In acknowledgement of the current inability to discern the source of the observed variance, many authors have begun to use combined terms such as “gender/sex” or “sex/gender” (Eliot et al., 2021; Franconi et al., 2019; Friedrichs & Kellmeyer, 2022; Hines, 2020; van Anders, 2022). Yet, despite this uncertainty, neuroscientific investigations often portray sex differences as “hardwiring” and imply a causal link between brain structure and genetic or hormonal sex differences (Friedrichs & Kellmeyer, 2022; Persson & Pownall, 2021). Therefore, more research is needed to disentangle the relationships between sex, gender, and brain structure.

Many researchers distinguish sex from gender according to biological versus sociocultural etiology, but this distinction is imperfect (Hyde et al., 2018). For instance, societal norms can affect how sex is interpreted or assigned (Carpenter, 2018; Griffiths, 2018), and gendered behavioral norms - such as those around diet, exercise, and sleep - can impact biology via epigenetics (Mauvais-Jarvis et al., 2020). To standardize research methods, the National Academies of Science, Engineering, and Medicine recently released a report on the measurement of sex and gender, defining sex as: “a multidimensional construct based on a cluster of anatomical and physiological traits that include external genitalia, secondary sex characteristics, gonads, chromosomes, and hormones.” Gender is defined as: “a multidimensional construct that links *gender identity*, which is a core element of a person’s individual identity; *gender expression*, which is how a person signals their gender to others through their behavior and appearance (such as hair style and clothing); and cultural expectations about social status, characteristics, and behavior that are associated with sex traits” (emphasis in original; National Academies of Sciences, 2022). Notably, these definitions eschew the typical binary categorizations of sex (male/female) and gender (masculine/feminine) and depart from normative views of gender as an immutable characteristic based on sex, as determined through physical examination (National Academies of Sciences, 2022). The shift in terminology reflects the increased visibility of transgender and intersex communities and the growing use of multi-dimensional gender assessments in research (Jacobson & Joel, 2018; Joel, 2011; Joel et al., 2014, 2015; Llaveria Caselles, 2021; Pelletier et al., 2015; Rippon et al., 2014). Despite these important efforts to discriminate sex- versus gender-related impacts in the biological sciences, few pediatric neuroscience studies to date have included both variables.

Many pediatric neuroimaging studies have reported sex differences in brain structure, albeit with little consensus on the location, magnitude, and potential functional relevance of such findings (DeCasien et al., 2022; Kaczkurkin et al., 2019; Raznahan et al., 2011). Subcortical brain regions such as the hippocampus, amygdala, and thalamus synthesize testosterone and estradiol (Ahmadpour & Grange-Messent, 2021; Azcoitia et al., 2011) and undergo volumetric changes during puberty (Vijayakumar et al., 2016). Nevertheless, meta-analyses of studies on hippocampus and amygdala volume have found no significant sex differences after normalizing for total brain volume (Marwha et al., 2017; A. Tan et al., 2016). Cortical thickness (cortical thickness) - the distance between the white matter boundary and the pial surface (Fischl & Dale, 2000) - decreases during child and adolescent development (Bethlehem et al., 2022), while regional surface area changes via decreases in gyrification index (GI) - the proportion of the gray matter surface located within sulcal folds (Alemán-Gómez et al., 2013; Schaer et al., 2012). Research on the role of sex on these cortical developmental outcomes has yielded conflicting results. Though some authors found sex differences in cortical thickness (Herting et al., 2015; Raznahan et al., 2016; Sepehrband et al., 2018) and gyrification (Fish et al., 2017; G. Li et al., 2014) others have reported no significant sex differences in children and adolescents (Alemán-Gómez et al., 2013; Mills et al., 2014; Vijayakumar et al., 2016; Wierenga et al., 2018). White matter microstructural differences, such as fractional anisotropy (FA) and mean diffusivity (MD), have also been extensively studied between the sexes during childhood. MD measures the average diffusion in three directions, while FA, on the other hand, describes the extent to which local brain structure restricts diffusion (O’Donnell & Westin, 2011). Males generally appear to have higher FA, but lower MD than females throughout the lifespan (Kaczkurkin et al., 2019; Raznahan et al., 2011), though some authors have found no adolescent sex differences in FA and MD (Giorgio et al., 2008, 2010; Lebel et al., 2008; Seunarine et al., 2016). Potential contrasting sex effects in pediatric neuroimaging studies could be due to small sample sizes, wide age ranges, or differences in sample composition. In this regard, the landmark Adolescent Brain Cognitive Development℠ Study (ABCD Study®), the largest long-term study of brain development in the United States (*ABCD Study*, 2022), enrolled over 11,000 9- and 10-year-old children to further study biological and social factors, like gender on brain development (Volkow et al., 2018). As such, several recent studies using ABCD data have reported significant neuroanatomical sex differences across numerous domains, including subcortical volume (Adeli et al., 2020), cortical thickness (Brennan et al., 2021; Tomasi & Volkow, 2023; Wiglesworth et al., 2023), gray matter density (Murray et al., 2022), white matter microstructure (Lawrence et al., 2021; Tomasi & Volkow, 2023), and differences in the association between brain structure and behavior (Chen et al., 2022; Kim et al., 2022). However, none of these previous studies on sex differences utilizing the ABCD study examined the potential role of gender.

In contrast to sex, gender has infrequently been used as a variable of interest in neuroscience research, let alone in studies focused on structural brain development. Articles mentioning gender often use the term interchangeably with sex, neglect to describe how gender was measured, and use binary, sex-related terminology, such as “male” and ”female”, making it difficult to evaluate (Backeljauw et al., 2015; Bhargava et al., 2021; Coviello et al., 2018; Fassett-Carman et al., 2022; Levman et al., 2017; Ou et al., 2015; Szulc-Lerch et al., 2018; Urben et al., 2017). This conflation exemplifies the lack of *a priori* model building in the existing neuroimaging literature on gender, which can lead to faulty assumptions of causality (Edmiston & Juster, 2022). For instance, when group differences in brain structure between transgender and cisgender participants are depicted as the result of atypical sexual differentiation (Kreukels & Guillamon, 2016; Kurth et al., 2022; Levin et al., 2022), it demonstrates both the assumption that sex *causes* any observed differences between sexes, and that gender identity is akin to brain sex. This reduces gender to a binary distinction between cisgender and transgender (Llaveria Caselles, 2021). In this dichotomy, studying the relationship between gender and brain structure in normative development is superfluous since sex and gender are assumed to be synonymous in the typically developing brain. It is therefore unsurprising that the handful of pediatric neuroimaging studies investigating gender and brain structure have largely focused exclusively on transgender children and utilized binary gender classification (Beking et al., 2020; Burke et al., 2014, 2016; Hoekzema et al., 2015; Nota et al., 2017; Skorska et al., 2021; Soleman et al., 2013; Staphorsius et al., 2015).

Moving beyond the cisgender/transgender dichotomy, **gender diversity** is used to describe variations of gender identity and expression that differ from those stereotypically associated with an individual’s sex assigned at birth (Bölte et al., 2023). Non-dichotomous assessments of both adults and children have demonstrated that gender diversity also exists among cisgender people (Joel et al., 2014; A. Potter et al., 2021). Yet, few pediatric neuroimaging studies to date have used a non-dichotomous measure of gender (Rauch & Eliot, 2022). Using the Child Sex Role Inventory, Wood *et al*. (2008) found femininity - but not masculinity - was associated with volume in portions of the frontal cortex (*n* = 74, ages 7.8-17.0) while Belfi *et al*. (2014), found masculinity - but not femininity - was associated with frontal and temporal cortex volume (*n* = 108, ages 7-17). In both studies, however, the results were only significant among female participants. Moreover, the Child Sex Role Inventory is adapted for children based on the Bem Sex Role Inventory, which was developed using ratings of the *desirability* of traits by 100 college students in the 1970s, making its contemporary validity questionable (Choi et al., 2008; Hoffman & Borders, 2001). Recently, Xerxa *et al*. asked participants whether they “wish to be the opposite sex” and whether they would “rather be treated as someone from the opposite sex?” with responses coded as ordinal variables as well as a binary categorization (e.g., adolescents who reported gender diversity and those who did not). While they found no association between gender diversity and gray matter volume, white matter volume, total brain volume, or regional subcortical volume, they did find that gender diverse male adolescents on average had thicker left inferior temporal cortices (Xerxa et al., 2023). Thus, despite more recent calls for increased attention to disentangling sex and gender in neuroscience research, their respective relationships with typical neurodevelopment remain equivocal. Sex differences studies consistently fail to account for gender as a possible confound, while neuroimaging studies of gender are rare, small, and concentrated on a subpopulation - limiting generalizability. Therefore, future research is needed to determine whether sex, gender, and/or their combination relate to neurodevelopmental outcomes, such as structural brain morphology.

To unify the previously separate lines of inquiry and address the common limitations of the existing sex and gender-based neuroimaging research, this cross-sectional study utilized data from the baseline ABCD Study visit, including multiple neuroimaging modalities, genetic data to assess X and Y allele frequency, and a continuous measure of gender. In order to establish clear, concise, and scientifically sound conceptions of sex and gender, we chose to limit our analysis to one dimension of sex - chromosomal sex - and one continuous measure of gender diversity that assesses how much a participant feels like a boy and how much they feel like a girl. The goal of the current study was to utilize a statistical modeling building approach to systematically examine sex and gender diversity as they relate to brain structure in children ages 9 and 10 years old. Specifically, we examined if sex, gender, or their combination captured the greatest amount of variance in common gray matter macrostructure outcomes, including cortical thickness, subcortical volumes, and gyrification, as well as white matter microstructure, including FA and MD. We hypothesized that we would be able to successfully capture variance uniquely associated with gender in the brain at ages 9-11 years old. Additionally, we anticipated that the best model for capturing outcome variance would differ by region and feature. In particular, since the few non-dichotomous gender studies to date have reported positive associations with the cortex and null findings with regard to the subcortex and global white matter volume, we expected gender to account for more regional variance in cortical architecture than in subcortical volume and white matter microstructure. Due to its documented association with sex hormones (Ahmadpour & Grange-Messent, 2021; Pletzer et al., 2018; Vijayakumar et al., 2021), we expected the subcortex to be primarily associated with sex alone.

## 2. Materials and Methods

### 2.1 Participants

Data for this analysis were acquired from the larger ongoing ABCD Study®. The Adolescent Brain Cognitive Development (ABCD) study enrolled 11,880 children from 21 different sites around the United States from 2016 through 2018 (*ABCD Study*, 2022; Casey et al., 2018; Hagler et al., 2019). Exclusion criteria for the ABCD study included lack of English proficiency, severe sensory, neurological, medical, or intellectual limitations, and inability to complete an MRI scan. The heterogeneity of sites captured geographic, demographic, and socioeconomic diversity of participants (Garavan et al., 2018; Heeringa & Berglund, 2020). In terms of age, sex, and household size, the ABCD cohort closely matches the distribution of 9- and 10-year-olds in the American Community Survey (ACS), a large probability sample survey of U.S. households conducted annually by the U.S. Bureau of Census (Heeringa & Berglund, 2020).

In the interest of reproducibility, the ABCD Study publicly shares its raw and processed data, as well as its processing pipelines. For the current study, we elected to use a combination of raw and tabulated questionnaire and neuroimaging data from the study baseline and one-year follow-up visit as obtained from the NDA 3.0 (raw T1 and T2 structural MRI files) and 4.0 (tabulated questionnaire and diffusion MRI) releases (NDA 3.0 and 4.0 data release 2021; https://dx.doi.org/10.15154/1523041). The rationale for performing our own preprocessing for gray matter macrostructure using raw T1w and T2w images, as compared to using the tabulated data for these outcomes, was that adding an additional contrast image improves pial surface parcellation (Lindroth et al., 2019; Misaki et al., 2015).

After obtaining the data, we performed a series of steps to determine the final sample for our analyses (Supplemental Figure 1). Participants were excluded if: data was collected outside the 21 primary research sites; incidental neurological findings were noted by a radiologist (Y. Li et al., 2021); participant was reported taking hormone medications; X and Y allele frequency ratio did not match their reported sex at birth; participant did not complete both felt-gender questions; quality control standards of the ABCD consortium for raw imaging or FreeSurfer reconstruction were not met (Hagler et al., 2019); or if errors occurred during execution of our pre- processing or processing pipelines. In addition, we elected to restrict our sample to one child per family to simplify the structure of our multilevel model and improve model fit. During the selection of a single child, to increase variability and improve our ability to assess associations between gender diversity and brain structure, we opted to include the child with the lowest felt-gender score from each family. In families where siblings had the same felt-gender score, one child was chosen randomly. Full description of the final sample for the current study can be found in Table 1.

**Table 1.** Sample Demographics.

|  | **ABCD**<br>**(N=11876)** | **Sample**<br>**(N=7693)** |
| --- | --- | --- |
| **Sex** |  |  |
| Female | 5680 (47.8%) | 3729 (48.5%) |
| Male | 6196 (52.2%) | 3964 (51.5%) |
| **Felt-Gender\*** |  |  |
| Mean (SD) | 4.74 (0.595) | 4.72 (0.612) |
| Median [Min, Max] | 5.00 [1.00, 5.00] | 5.00 [1.00, 5.00] |
| Missing | 840 (7.1%) | 0 (0%) |
| **Age (months)** |  |  |
| Mean (SD) | 119 (7.50) | 119 (7.46) |
| Median [Min, Max] | 119 [107, 133] | 119 [107, 133] |
| **Pubertal Development** |  |  |
| Pre | 5837 (49.1%) | 3805 (49.5%) |
| Early | 2713 (22.8%) | 1765 (22.9%) |
| Mid/Late | 2854 (24.0%) | 1849 (24.0%) |
| Missing | 472 (4.0%) | 274 (3.6%) |
| **Race\*** |  |  |
| White | 7517 (63.3%) | 5009 (65.1%) |
| Black | 1868 (15.7%) | 1060 (13.8%) |
| Multiracial (Black) | 649 (5.5%) | 401 (5.2%) |
| Multiracial (Non-black) | 785 (6.6%) | 508 (6.6%) |
| Other<sup>a</sup> | 874 (7.4%) | 597 (7.8%) |
| Missing | 183 (1.5%) | 118 (1.5%) |
| **Ethnicity** |  |  |
| Non-Hispanic | 9312 (78.4%) | 5989 (77.9%) |
| Hispanic | 2411 (20.3%) | 1611 (20.9%) |
| Missing | 153 (1.3%) | 93 (1.2%) |
| **Parent Education\*** |  |  |
| < HS Diploma | 578 (4.9%) | 334 (4.3%) |
| HS or GED | 1110 (9.3%) | 643 (8.4%) |
| Some College | 3058 (25.7%) | 1922 (25.0%) |
| Bachelor | 3010 (25.3%) | 1963 (25.5%) |
| Post Graduate Degree | 4041 (34.0%) | 2777 (36.1%) |
| Missing | 79 (0.7%) | 54 (0.7%) |

\* Difference between research sample and ABCD Study sample is statistically significant at the $p < 0.05$ level.
<sup>a</sup> The "Other" race/ethnicity category includes participants who were parent-identified as Asian Indian, Chinese, Filipino/a, Japanese, Korean, Vietnamese, Other Asian, American Indian/Native American, Alaska Native, Native Hawaiian, Guamanian, Samoan, Other Pacific Islander, or Other Race

After performing stringent data cleaning and removing siblings, our final sample still closely matched the ABCD sample in terms of sex, age, pubertal development, and ethnicity, but differed significantly in terms of race and parental education. While these differences do not impact the internal validity of the study, it could impact the generalizability of our results. To ensure our purposeful selection to increase greater gender diversity did not confound our results, we compared our final analytic sample to one in which one child per family was selected at random (rather than by felt-gender score), which showed similar distributions of all other covariates included in our models. This suggests the other factors in subset selection (i.e., image quality (Cosgrove et al., 2022)) possibly led to sample differences rather than the oversampling for gender diversity.

### 2.2 Neuroimaging Data

A harmonized data collection protocol was utilized across sites with either a Siemens, Phillips, or GE 3T MRI scanner. Motion compliance training, as well as real-time, prospective motion correction was used to reduce motion distortion. T1w images were acquired using a magnetization-prepared rapid acquisition gradient echo (MPRAGE) sequence (TR=2500, TE=2.88, flip angle=8) and T2w images were obtained with fast spin echo sequence (TR=3200, TE=565, variable flip angle), with 176 slices with 1mm^3^ isotropic resolution (Casey et al., 2018). Diffusion MRI data was acquired in the axial plane at 1.7 mm3 isotropic resolution with multiband acceleration factor 3. Ninety-six non-collinear gradient directions were collected with seven b0 images and four non-zero b-values (b=500, b=1000, b=2000, b=3000). Trained technicians inspected all T1w, T2w, and dMRI images and all images underwent a centralized quality control process in order to identify severe artifacts or irregularities (Hagler et al., 2019).

To assess gray matter macrostructure, we obtained the raw baseline T1w and T2w images from the ABCD 3.0 release (NDA 3.0 data release 2020; https://dx.doi.org/10.15154/1520591) and implemented the Human Connectome Project minimal preprocessing pipeline (Glasser et al., 2013) at the Stevens Institute of Neuroimaging and Informatics. Regional parcellation and segmentation was then performed based on the Desikan-Killiany atlas using FreeSurfer 7.1.1 for each participant using T1w and T2w images (Desikan et al., 2006). Calculation of lGI was accomplished using the additional FreeSurfer function “lgi” (Schaer et al., 2012), which compares total surface area within sulcal folds to visible pial surface area. This function is sensitive to topological defects in the FreeSurfer parcellations; therefore, in cases where lGI calculation failed due to aberrant results, participants were excluded from all analyses. Primary outcomes of interest included 68 cortical thickness regions, 22 subcortical volume regions, 68 lGI regions, 21 white matter regions for FA and MD. For a complete list of regions by feature, please see Supplemental Table 3.

Tabulated white matter microstructure results and demographic data from the baseline study visit were obtained from the 4.0 data release via the NIMH Data Archive (https://nda.nih.gov/abcd/; http://dx.doi.org/10.15154/1523041). ABCD diffusion processing employs five iterations of eddy current correction and robust tensor fitting to minimize gradient distortions and motion (Hagler et al., 2019; Hagler Jr et al., 2009). The b=0 images are coarsely registered to a diffusion atlas before being registered to T1w images using mutual information. DMRI images are then resampled and registered using the transform from rigid registration of the T1w image to the diffusion atlas. Finally, the diffusion gradient matrix is adjusted for head rotation. Probabilistic atlas-based tractography is then performed with AtlasTrack using *a priori* tract location probabilities to inform fiber selection (Hagler Jr et al., 2009). For this study, we utilized the tabulated FA and MD data from the AtlasTrack fiber atlas. Specifically, we selected the fornix, cingulate cingulum, parahippocampal cingulum, uncinate fasciculus, superior longitudinal fasciculus, inferior longitudinal fasciculus, inferior fronto-occipital fasciculus, anterior thalamic radiations, corticospinal tracts, forceps major, forceps minor, and corpus callosum as regions of interest (ROIs).

### 2.3 Independent variables

#### 2.3.1 Sex

The ABCD Study collects parent-reported sex assigned at birth. However, due to the multidimensional nature of sex, assignment at birth is not always an accurate reflection of chromosomal sex. Therefore, we also chose to use the frequency ratio of X and Y alleles from blood samples to detect the presence of a Y chromosome and ascertain the genetic sex of participants. Children whose assigned sex and genetic sex did not match (n = 9) were excluded from analysis. Furthermore, children who were reported to be taking hormone medications at the baseline assessment were excluded from analysis (for a list of excluded medications, see Supplemental Table 1).

#### 2.3.2 Gender Diversity

At the one-year visit, children completed a gender questionnaire (A. Potter et al., 2021; A. S. Potter et al., 2022). The first two questions of the questionnaire asked how much the participant feels like a boy and how much they feel like a girl, on a 5-point Likert scale (“Not at all,” “A little,” “Somewhat,” “Mostly,” “Totally”). Scoring was based on sex assigned at birth, with questions reverse-coded for the gender that was culturally incongruent with the participant’s sex assigned at birth (e.g., how much an XX child feels like a boy and how much an XY child feel like a girl) (see Supplemental Table 2). Then the two scores were averaged to create a “felt-gender” score which represented the level of congruence between the participant’s personal feelings of gender and the sex/gender assigned at birth). In the current study, we used the term “gender diversity” to refer to deviation from the maximum felt-gender score of 5 (e.g., feeling “totally” congruent with the gender assigned to one’s birth sex and “not at all” having incongruent gender feelings). As such, lower felt-gender scores reflect higher levels of gender diversity.

#### 2.3.3 Other Predictors

Based on our questions of interest, we identified a number of demographic variables in the ABCD dataset as important predictors of brain structure. **Age** at scan date was measured in months and rounded to the nearest whole month. **PDS** was calculated based on the parent-report version of the Pubertal Development Scale and categorized as prepuberty, early puberty, mid puberty, late puberty, and post-puberty (Cheng et al., 2021; Herting et al., 2021; Petersen et al., 1988; Thijssen et al., 2020). Since few children in this age range were in late puberty or post-pubertal, we combined the mid, late, and post-puberty groups into a single category (mid/late puberty). We chose to include measures of **race**, **ethnicity**, and **socioeconomic status** in our models because human neurodevelopment is sensitive to myriad ecological factors which, due to systemic social injustice, are correlated with sociocultural variables, like race, ethnicity, and socioeconomic status (Nketia et al., 2021; Werchan & Amso, 2017). **Race** was determined via parent report, with parents encouraged to select all answers that apply. For participants who selected more than one race, we categorized them as multiracial Black if one of their selections was “Black” or multiracial non-Black. We combined the categories of Asian Indian, Chinese, Filipino/a, Japanese, Korean, Vietnamese, Other Asian, American Indian/Native American, Alaska Native, Native Hawaiian, Guamanian, Samoan, other Pacific Islander, and “other race” into a single category (“other race”), due to small group numbers. **Ethnicity** was also parent-reported as either Hispanic or non-Hispanic. Multiple measurements of socioeconomic status - such as parental educational attainment, household income, income-to-needs ratio, neighborhood disadvantage - have been linked to brain structure in studies of ABCD data (Rakesh et al., 2022). Due to its low non-response rate, we chose to use **educational attainment**, which was defined as the highest level of education achieved by any parent in the household and binned into the following categories: less than high school diploma, high school diploma or GED, some college, bachelor’s degree, or postgraduate degree. **Scanner** manufacturer was included to account for differences in data collection caused by MRI software and hardware. Although intracranial volume is often used account for differences in brain size during brain volume analyses, actual brain volume varies throughout the course of the day (Karch et al., 2019; Nakamura et al., 2015; Trefler et al., 2016). Thus, we chose to also include **total brain volume (TBV)** as a predictor in our subcortical volume models to account for the relationship between regional and whole-brain volume. TBV was calculated by FreeSurfer, then scaled by the sample root-mean-square.

### 2.4 Analyses

Mixed effects modeling is a statistical approach for handling nested data structures, such as in the case of multi-site studies that need to account for site clustering that may result from unmeasured sociocultural or environmental factors of participants and/or the data collection process (Heeringa & Berglund, 2020; Owens et al., 2021). The term “mixed” refers to the fact that it combines traditional fixed effect modeling with addition of random effects - variables whose effect on the outcome is not uniform across groups (Kreft & De Leeuw, 1998). Our study implemented the lmer() and glmer() functions for generalized linear mixed modeling from the lme4 package in R (Bates et al., 2015; R Core Team, 2019) to examine the effects of sex, felt-gender and their interaction on gray matter macrostructure and white matter microstructure. Residuals from all models were visually and numerically inspected to rule out violations of model assumptions. If a model failed to meet distributional assumptions (i.e., normality assumption) and/or experienced convergence issues with lmer(), we implemented glmer() to fit the model with gamma distribution assumption but keeping the identity link to preserve interpretability. In order to improve the accuracy of our models, we sought to account for additional variables that influence neurodevelopment, and to account for nesting of subjects within sites, we included the data collection **site** as a random effect. For our fixed effects, we added age, PDS, race, ethnicity, parental education, and scanner as fixed effects (all collected at the baseline visit), as well as TBV for the models of subcortical volume.

For this study, we relied on a model building strategy to inform our understanding of how sex, gender, and their combination (i.e., main effects and/or interaction) relate to gray matter macrostructure and white matter microstructure. For each ROI outcome, we fit a null model without sex or gender, along with four models of sex and/or gender. The null model (M0) included only our covariates (e.g., ROI ∼ age + PDS score + race + ethnicity + parental education + scanner + (1 | site)) (and when applicable, TBV for subcortical outcomes). Next, sex was added to the M0 model as a fixed effect to create our sex model (M1), whereas felt-gender score was added to the M0 model as a fixed effect to create a gender model (M2). Next, we created two additional models that included both sex and gender as main effects in a sex and gender model (M3), as well a fourth model that included both main effects and a sex-by-gender interaction term for a sex-by-gender model (M4). The point biserial correlation between sex and felt-gender scores in our analytic sample was -0.24. To rule out multicollinearity amongst our covariates, we calculated the variance inflation ratio (vif) of each model. Importantly, all variance inflation factor values were below 3, suggesting no multicollinearity between variables in any model. Given that very large samples tend to transform small differences into statistically significant effects, we chose to calculate Cohen’s f^2^ to quantify the effect size and aid in our interpretation of the regional relationships detected between structure and sex, gender, and interaction.

For each ROI, we then used pairwise ANOVA tests in order to determine whether each model captured more variance than the null model as well as to compare Akaike Information Criterion (AIC) between models with differing numbers of independent variables (see Supplemental Figure 2). In cases where no model fit the outcome variance better than the null model (M0), the null model was considered the best fit for the data. If the interaction model (M4) had a p-value < 0.05 in pairwise ANOVA comparisons with all three other models (M1, M2, and M3) and the null model (M0), it was chosen as the best model. Otherwise, if the sex + gender model (M3) had a p-value < 0.05 in pairwise comparisons with the sex model (M1), the gender model (M2), and the null model (M0) it was chosen as the best model. If neither the interaction nor the sex + gender model was chosen as the best and only one of the two remaining models - sex (M1) and gender (M2) - significantly improved fit compared to the null model, then the model that performed better than the null was considered the best-fitting model. If both the sex (M1) and gender (M2) models improved fit compared to the null model, then the model with the lowest AIC was chosen as the best model for explaining outcome variance. Finally, to gauge the magnitude of the difference in model fit, we calculated the range of model *r^2^* values for each ROI.

## 3. Results

Below, we report model comparison results by macrostructural (i.e., subcortical volume, cortical thickness, and gyrification index) and microstructural outcomes of interest, respectively. A summary of which model was deemed to account for the greatest amount of variance by macrostructural gray matter outcomes is presented in Table 2 (as well as in Supplemental Tables 5, 6, & 7), whereas white matter microstructural outcomes are presented in Table 3 (as well as Supplemental Tables 8-9). In the majority of both gray matter and white matter ROIs, structural variance was best accounted for by the sex model (M1). The gender model (M2) was not the best model for capturing structural variance in any white or gray matter regions; however, the sex and gender model (M3) captured more variance than sex alone in eight gray matter regions and the interaction model (M4) captured the most variance in one white matter region.

**Table 2:** Number and frequency (%) of the best model for each gray matter macrostructural ROI in terms of outcome variance captured (R^2^). Abbreviations: M0 = null model (i.e., covariates only); M1 = null model + sex; M2 = null model + gender; M3 = null model + sex + gender; M4 = null model + sex + gender + sex-by-gender interaction; ROI = regions of interest.

| Gray Matter | Total ROIs | Model |  |  |  |  |
| --- | --- | --- | --- | --- | --- | --- |
|  |  | M0 | M1 | M2 | M3 | M4 |
|  | 158 | 31 (19.6%) | 119 (75.3%) | 0 (0.0%) | 8 (5.1%) | 0 (0.0%) |
| **Subcortical Volume** | **22** | **5 (22.7%)** | **15 (68.2%)** | - | **2 (9.1%)** | - |
| Basal Ganglia | 8 | 1 (12.5%) | 7 (87.5%) | - | - | - |
| Limbic | 6 | 4 (66.7%) | 2 (33.3%) | - | - | - |
| Corpus Callosum | 5 | - | 5 (100%) | - | - | - |
| Other | 3 | - | 1 (33.3%) | - | 2 (66.7%) | - |
| **Cortical Thickness** | **68** | **23 (33.8%)** | **44 (64.7%)** | - | **1 (1.5%)** | - |
| Frontal | 22 | 8 (36.4%) | 13 (59.1%) | - | 1 (4.5%) | - |
| Temporal | 18 | 3 (16.7%) | 15 (83.3%) | - | - | - |
| Parietal | 10 | 6 (60.0%) | 4 (40.0%) | - | - | - |
| Occipital | 8 | 4 (50.0%) | 4 (50.0%) | - | - | - |
| Cingulate | 8 | 2 (25.0%) | 6 (75.0%) | - | - | - |
| Insula | 2 | - | 2 (100%) | - | - | - |
| **Local Gyrfication Index** | **68** | **3 (4.4%)** | **60 (88.2%)** | - | **5 (7.4%)** | - |
| Frontal | 22 | 3 (13.6%) | 18 (81.8%) | - | 1 (4.5%) | - |
| Temporal | 18 | - | 17 (94.4%) | - | 1 (5.6%) | - |
| Parietal | 10 | - | 7 (70.0%) | - | 3 (30.0%) | - |
| Occipital | 8 | - | 8 (100%) | - | - | - |
| Cingulate | 8 | - | 8 (100%) | - | - | - |
| Insula | 2 | - | 2 (100%) | - | - | - |

**Table 3:** Number and frequency (%) of the best model for each white matter microstructural ROI in terms of variance explained. Abbreviations: M0 = null model (i.e., covariates only); M1 = null model + sex; M2 = null model + gender; M3 = null model + sex + gender; M4 = null model + sex + gender + sex-by-gender interaction; ROI = regions of interest.

| White Matter | Total ROIs | Model |  |  |  |  |
| --- | --- | --- | --- | --- | --- | --- |
|  |  | M0 | M1 | M2 | M3 | M4 |
|  | 42 | 8 (19.1%) | 33 (78.6%) | 0 (0%) | 0 (0%) | 1 (2.4%) |
| **Fractional Anisotropy** | **21** | **6 (28.6%)** | **15 (71.4%)** | - | - | - |
| Association Fibers | 6 | - | 6 (100%) | - | - | - |
| Projection Fibers | 4 | 2 (50.0%) | 2 (50.0%) | - | - | - |
| Limbic Fibers | 8 | 3 (37.5%) | 5 (62.5%) | - | - | - |
| Callosal Fibers | 3 | 1 (33.3%) | 2 (66.7%) | - | - | - |
| **Mean Diffusivity** | **21** | **2 (9.5%)** | **18 (85.7%)** | - | - | **1 (4.8%)** |
| Association Fibers | 6 | 1 (16.7%) | 5 (83.3%) | - | - | - |
| Projection Fibers | 4 | - | 4 (100%) | - | - | - |
| Limbic Fibers | 8 | - | 7 (87.5%) | - | - | 1 (12.5%) |
| Callosal Fibers | 3 | 1 (33.3%) | 2 (66.7%) | - | - | - |

### 3.1 Gray Matter Macrostructure

#### 3.1.1 Subcortical Volume

All subcortical volume averages were larger in males than in females. Using the modeling approach to determine which factors best explained the variance subcortical volume, we found that in the majority of subcortical ROIs (15/22; 68%), the sex model (M1) captured more variance than the other models (Figure 1). However, the direction of the relationship between sex and volume varied by region (Figure 2). The model with both sex and gender (M3) was determined to best capture the variance in the left and right cerebellum; albeit sex was a significant predictor in these models, while gender diversity was not statistically significant. The gender model (M2) and the interaction model (M3) were not the best-fitting model for any subcortical region. In the left and right thalamus, left and right hippocampus, and right pallidum, neither the sex model (M1), the gender model (M2), the sex + gender model (M3), nor the interaction model (M4) explained more variance than the null model; therefore we consider the null model (M0; covariates only) to be the best-fitting model in 23% of subcortical regions examined, including the right pallidum, bilateral hippocampus, and bilateral thalamus. Effect sizes of individual sex, gender, and interaction beta coefficients in each model were small (max *f^2^* = 0.028). Additionally, the difference in *R^2^* between the best model and the null model was universally small (mean Δ*R^2^* = 0.004, max Δ*R^2^* = 0.014).

**Figure 1.**
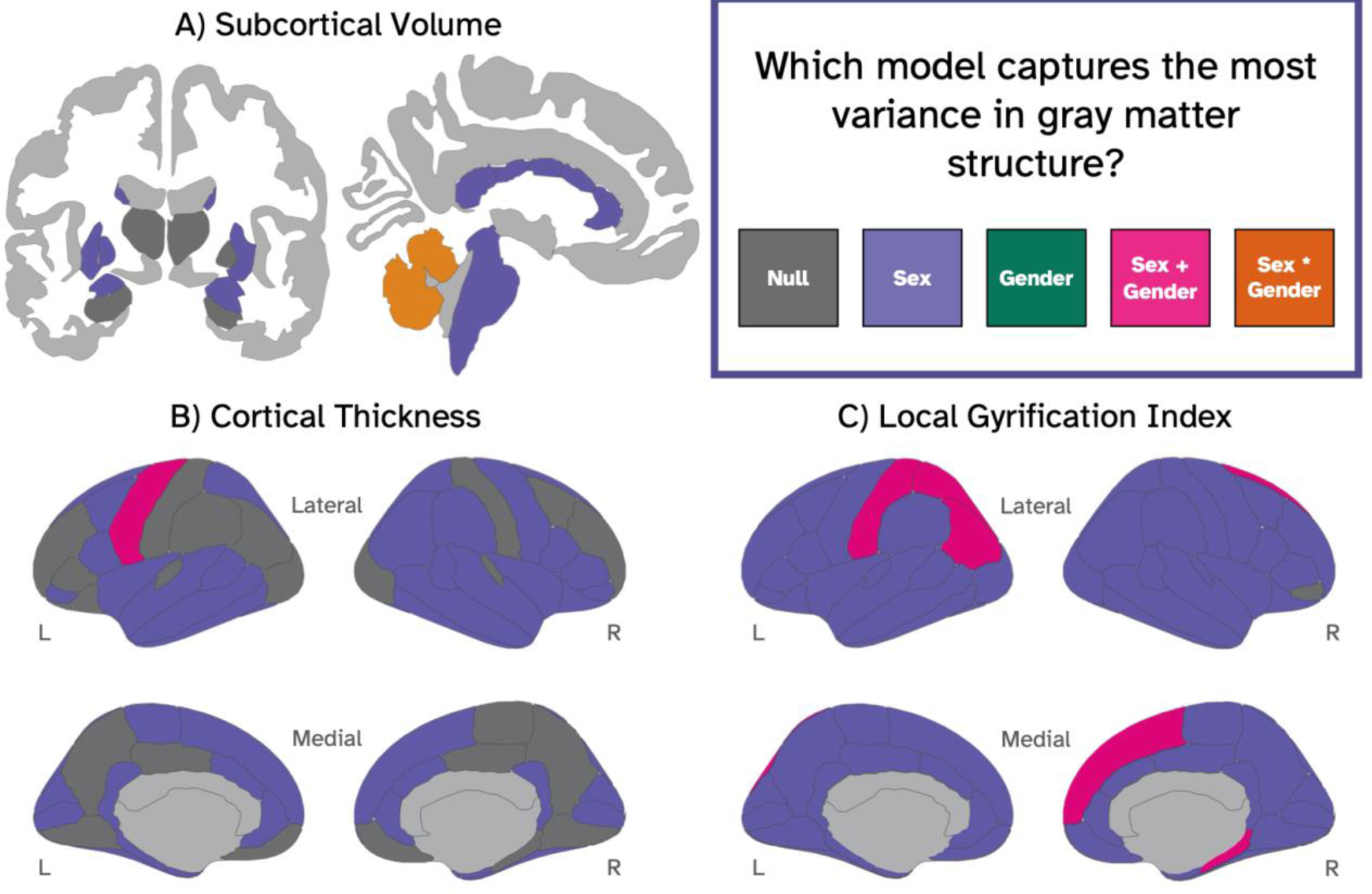
Coronal and sagittal slices depicting the best model for capturing variance in gray matter structure, including subcortical volume (A), cortical thickness (B), and local gyrification index (C). Each ROI is colored according to the best-fitting model: gray = null model (i.e., covariates only); purple= M1 (null model + sex; green = M2 (null model + gender); pink = M3 = (null model + sex + gender); orange= M4 (null model + sex + gender + sex-by-gender interaction).

**Figure 2.**
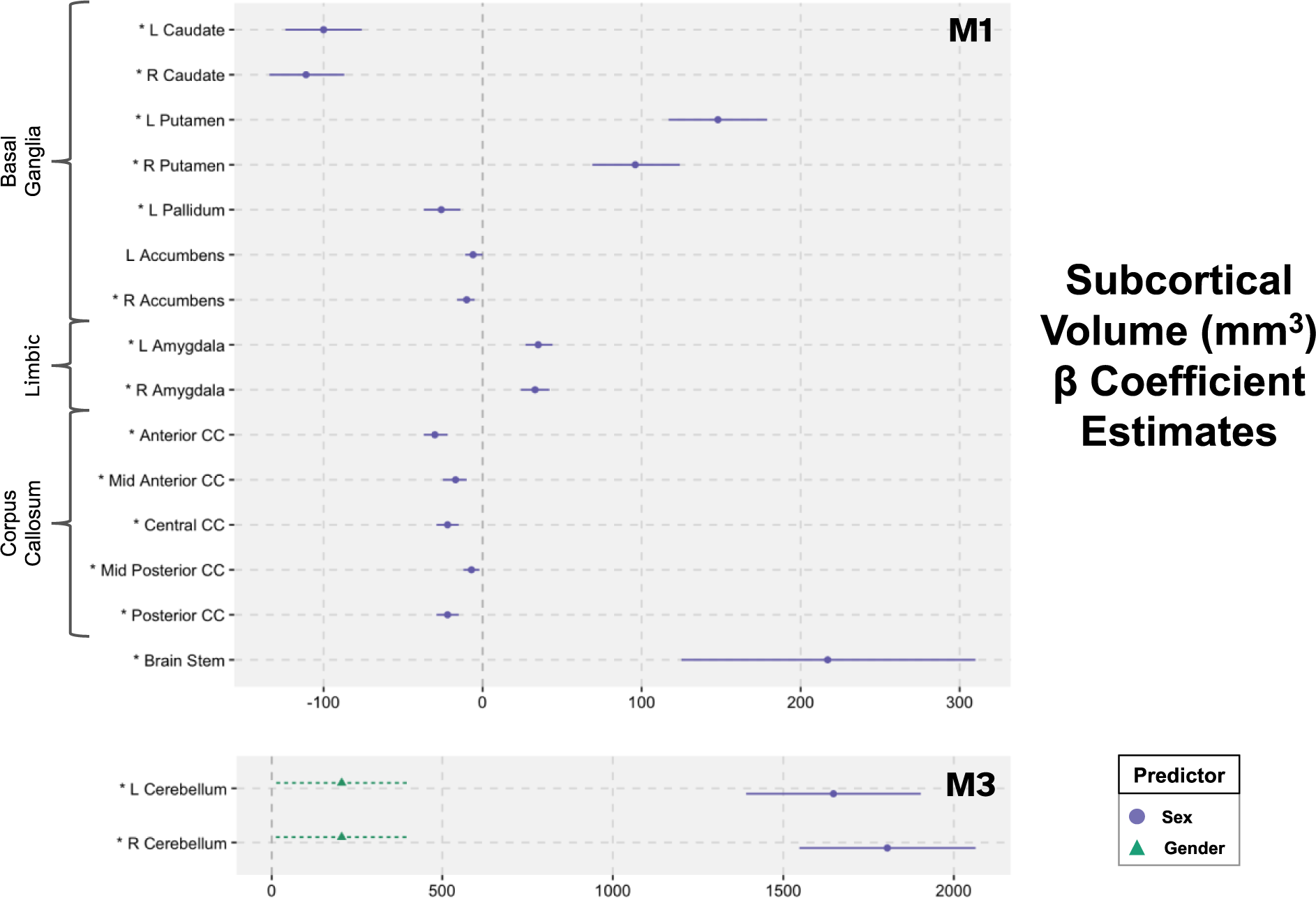
Beta coefficient estimates and 95% confidence intervals (CI) for the best-fitting model for each subcortical region. Abbreviations: M1 = null model (covariates) + sex; M3 = null model (covariates) + sex + gender; R=right; L=left; * = FDR- corrected p-value for sex < 0.05

#### 3.1.2 Cortical Thickness

With regard to variance in regional cortical thickness, we found that the sex model (M1) captured the most variance in 65% (44/68) of ROIs examined (Figure 1). The model with a combination of sex and gender (M3) was the best fit for the variance in thickness of the left precentral gyrus. In this region, males showed thinner cortex than females (β = -0.012, *p* < 0.001), while the association between gender and thickness was not significant. For 34% of regions, none of the models performed better than the null model (M0; covariates only) - including bilaterally in the rostral middle frontal gyrus, medial orbitofrontal cortex, transverse temporal gyrus, postcentral gyrus, precuneus, lateral occipital cortex, lingual gyrus, and posterior cingulate cortices, the null model was the best model bilaterally. Where significant, the relationship between sex and cortical thickness was heterogeneous (see Figure 3): in the left middle temporal, left and right fusiform gyrus, left parahippocampal gyrus, left and right superior parietal lobe, and right caudal anterior cingulate cortex, the mean cortical thickness among males was higher than the mean among females; in the other 34 regions where the difference between sexes was significant, the mean cortical thickness measurement for females was higher than the mean for males.

**Figure 3.**
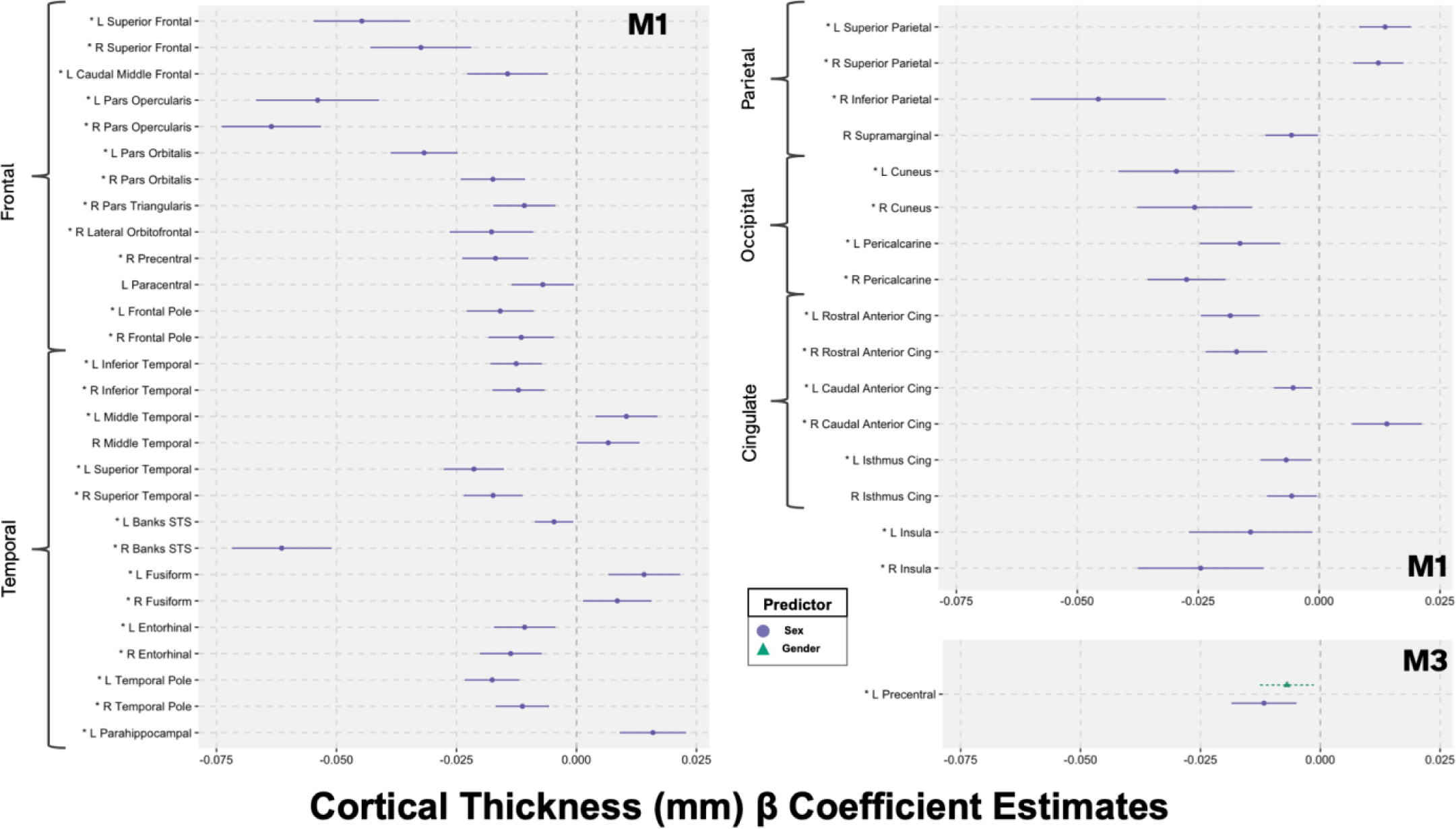
Beta coefficient estimates and 95% confidence intervals (CI) for the best-fitting model for each cortical thickness region. Abbreviations: M1 = null model (covariates); M3 = null model (covariates) + sex + gender; R=right; L=left; Cing = cingulate; * = FDR-corrected p-value for sex < 0.05

The overlap between the male and female confidence intervals was large in every region. Across all models and ROIs, the effect sizes of our sex and gender estimates were very small (max *f^2^* = 0.020), and the difference in explained variance between models was minimal (as measured with conditional *R^2^* range per ROI; mean Δ*R^2^* = 0.002, max Δ*R^2^* = 0.018).

#### 3.1.3 Local Gyrification Index

The sex model (M1) best described the variance in gyrification in 88% of regions, while the sex + gender model (M3) was the best predictor of variance in the right superior frontal cortex, right parahippocampal gyrus, left superior parietal cortex, left inferior parietal cortex, and left postcentral gyrus (Figure 1). In all 68 ROIs, the mean local gyrification index among males was higher than the mean among females (Figure 4). The association between felt-gender score and lGI was not significant in any region. Sex and gender estimate effect sizes were again small according to Cohen’s guidelines (Cohen, 1988) for interpretation of effect size (max *f^2^* = 0.077).

**Figure 4.**
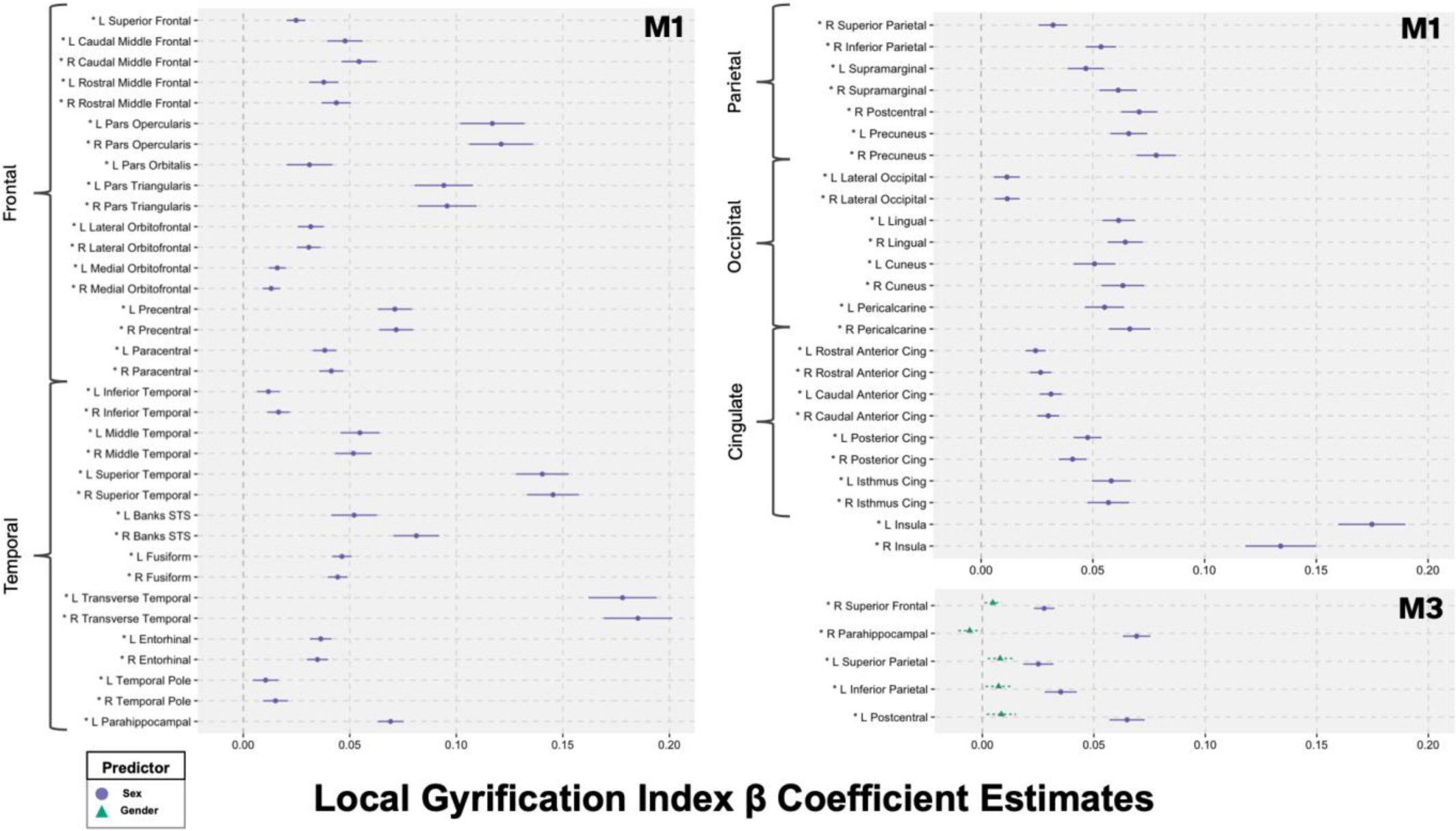
Beta coefficient estimates and 95% confidence intervals (CI) for the best-fitting model for local gyrification of each region. Abbreviations: M1 = null model (covariates) + sex; M3 = null model (covariates) + sex + gender; R=right; L=left; * = FDR-corrected p-value for sex < 0.05

However, the maximum effect size for lGI - which was observed in the right superior temporal gyrus - was also the maximum effect size in this study, suggesting that sex accounts for a larger proportion of the variance in lGI than it does in any other metric examined. Furthermore, lGI outcomes had the largest average difference in explained variance between the best and worst model (mean Δ*R^2^* = 0.023, max Δ*R^2^* = 0.064).

### 3.2 White Matter Microstructure

#### 3.2.1 Fractional Anisotropy

Model building results showed that mean FA was best explained by the sex model (M1) in 71% of tracts (15/21; Figure 5). In half of fiber tracts, the male FA average was higher than the female average, while the female average was higher than males in the other half (Figure 6). For 29% of regions (6/21), none of the models accounted for significantly more variance than the null model with covariates (M0), including the forceps major, left uncinate fasciculus, bilateral superior longitudinal fasciculus, and bilateral corticospinal tracts. Despite the sex model accounting for the most variance of the four models, only a small proportion of FA variance was attributable to sex (max *f^2^* = 0.017) and the difference between models in terms of explained variance was small (mean Δ*R^2^* = 0.003, max Δ*R^2^* = 0.012).

**Figure 5.**
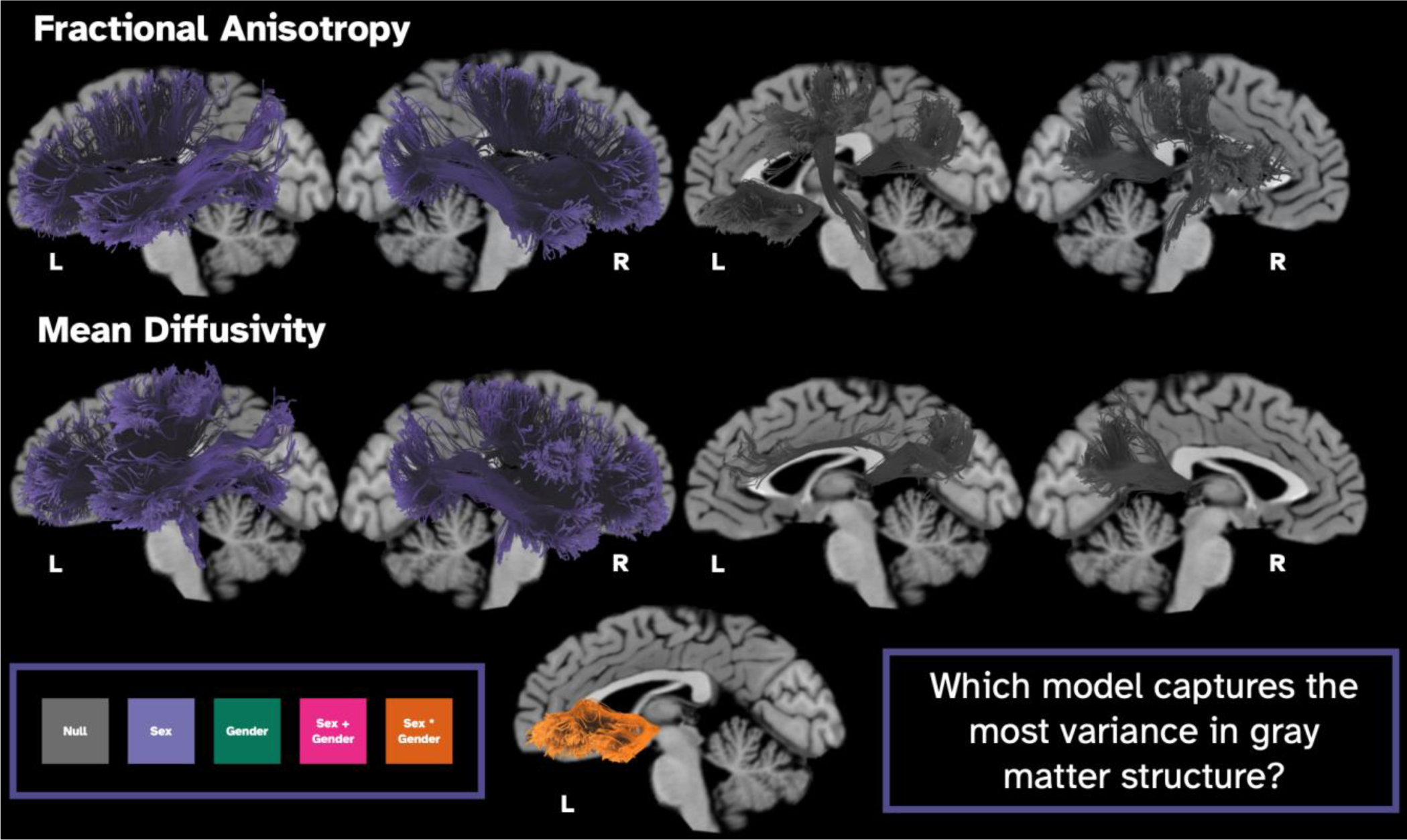
T1-weighted images overlaid with examined white matter tracts colored according to the best-fitting model. White = null model (i.e., covariates only); purple= M1 null model (covariates) + sex; green = M2 (null model + gender); pink = M3 (null model + sex + gender); orange= M4 (null model + sex + gender + sex-by-gender interaction). The gender model (M2) and the sex + gender model (M3) were not the best-fitting model in any WM ROIs.

**Figure 6.**
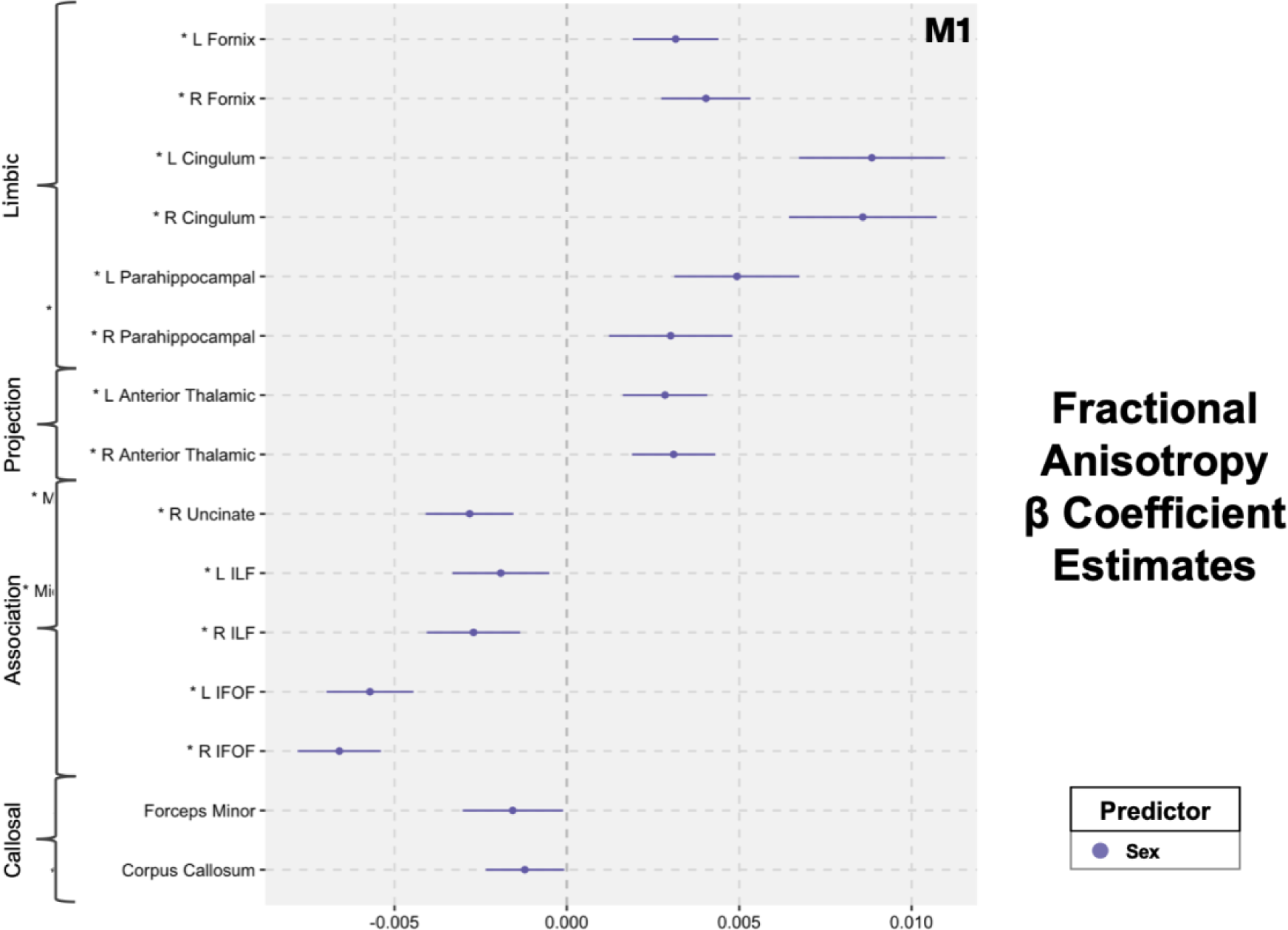
Beta coefficient estimates and 95% confidence intervals (CI) for the best-fitting model for FA in each white matter tract. Abbreviations: M1 = null model (covariates) + sex; R=right; L=left; ILF = inferior longitudinal fasciculus; IFOF = inferior fronto-occipital fasciculus

#### 3.2.2 Mean Diffusivity

In modeling analyses, the sex model (M1) best described the variance in MD in 86% of regions (18/21) including limbic, projection, association, and callosal white matter fiber tracts (Figure 5). The interaction model (M4) best accounted for MD variability in the left uncinate, and sex - but not gender diversity or the interaction term - was independently significantly associated with MD in the model. In 10% of tracts (2/21), MD variance was best explained by the null model (M0), including the left cingulate cingulum, and forceps major. In all regions where sex was a significant predictor of MD, the male average MD was higher than the female average (Figure 7). Effect sizes were small (max *f^2^*= 0.035), and so was the difference in variance explained by each model, as measured by conditional *R^2^* range (mean Δ*R^2^* = 0.008, max Δ*R^2^* = 0.026).

**Figure 7.**
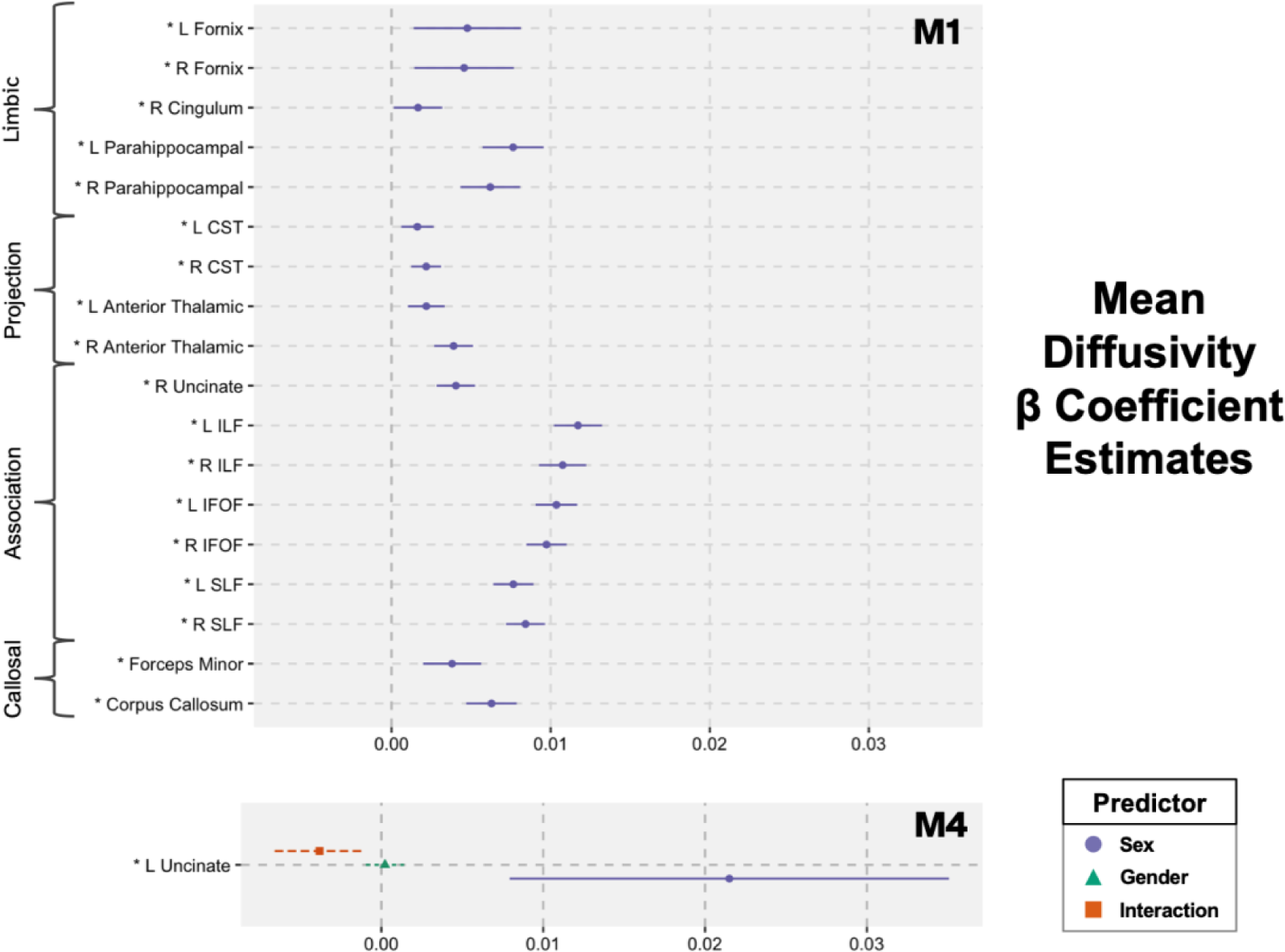
Beta coefficient estimates and 95% confidence intervals (CI) for the best-fitting model for MD in each white matter tract. Abbreviations: M1 = null model (covariates) + sex; M4 = null model (covariates) + sex + gender + sex-by-gender interaction); R=right; L=left; CST = corticospinal tracts; ILF = inferior longitudinal fasciculus; IFOF = inferior fronto-occipital fasciculus; SLF = superior longitudinal fasciculus

## 4. Discussion

This study is the largest neuroimaging study to date using a non-dichotomous assessment of gender (Rauch & Eliot, 2022), and the first to examine the relationship between diversity of felt–gender and diversity of brain structure - while also considering sex - in a nationwide pediatric sample. The classical model (M1) - with sex (but not gender) - explained the most variance in 75% of gray matter ROIs and 79% of white matter ROIs. This suggests a widespread association between sex and both gray and white matter structures in youth ages 9-11 years-old - although the effect sizes for sex estimates were small in all cases. In contrast, the gender model (M2) did not account for more variance than the other models in any of the regions examined. The sex + gender model (M3) captured the most variance in 8 gray matter ROIs (left and right cerebellum volume, left precentral cortical thickness, and local gyrification index in the left superior and inferior parietal cortex, left precentral gyrus, and right parahippocampal gyrus), while the interaction model (M4) captured the most variance in left uncinate fasciculus MD. Nonetheless, the β coefficient estimates for gender and the sex-by-gender interaction term from the M4 model were not statistically significant in the best model for any region. Taken together, these results demonstrate that, in 9-11-year-olds, felt-gender diversity is not directly associated with brain structure, while sex accounts for a small amount of variance in brain structure across a wide array of regions and measurements.

Our findings align with previous work involving the ABCD dataset where different types of machine learning techniques were implemented to probe potential discriminative sex differences in the adolescent brain (Adeli et al., 2020; Brennan et al., 2021; Kim et al., 2021, 2022). For example, deep learning identified the subcortex as the most sex-discriminative area of the brain (Adeli et al., 2020). The authors further probed the ROIs associated with the most-discriminative voxels using two-sample *t*-tests and reported sex differences in volume in the pallidum, some parts of the cerebellum, hippocampus, and putamen - but not the amygdala or insula (Adeli et al., 2020). In another ABCD analysis, machine learning analysis utilizing various brain features predicted sex with 70.8% accuracy with as few as 13 attributes (select ROIs pertaining to cortical thickness, cortical volume, sulcal depth, gray matter intensity, white matter intensity, and gray/white intensity contrast)(Brennan et al., 2021).

Moreover, these analyses did not account for other possible sources of variance between males and females, such as pubertal development, which differs significantly between males and females at ages 9-11 years in the ABCD sample (Herting et al., 2021), nor did they consider the potential role of gender.

Through our model building approach and adjusting for these key covariates, our analysis supported previous findings of significant sex differences in both gray and white matter structures as well as highlighted a number of brain regions that are similar between the sexes (i.e., best explained by the null model) at ages 9-11 years old. Similar to Adeli and colleagues (Adeli et al., 2020), we confirmed significant sex differences bilaterally in the caudate, putamen, pallidum, cerebellum. We found additional sex differences in the brain stem, corpus callosum, nucleus accumbens, and the amygdala. Experiments in mice using the “Four Core Genotypes” model (XX and XY gonadal males, and XX and XY gonadal females) demonstrate that the X chromosome and gonadal hormones influence the volume of the cerebellum and corpus callosum, providing a potential biological explanation for our findings (Corre et al., 2016). In contrast to Adeli, we found no significant association between sex and hippocampal volume in any model, despite the known role of the hippocampus in the synthesis of both testosterone and estradiol (Ahmadpour & Grange-Messent, 2021; Azcoitia et al., 2011) as well as its enrichment in estrogen receptors (Bean et al., 2014). Research in murine models of sex chromosome aneuploidies has found that, rather than increasing differences, X-chromosome gene dosage effects and androgen effects counteract each other in some brain regions and reduce sex differences in volume, (Raznahan et al., 2013). This demonstrates that a regional association with sex hormones does not necessitate a sex difference in structure. Instead, sex interacts with neuroanatomy through multiple mechanisms, including gene expression, X-chromosome inactivation and inactivation escape, endocrine regulation, and receptor density (Raznahan & Disteche, 2021; Sigurdardottir et al., 2020). The relationship between cortical thickness and sex is also complex. Cortical thickness does not appear to be related to X-chromosome inactivation (Mallard et al., 2021), and findings regarding its association with androgen levels in children are mixed (Herting et al., 2015; Nguyen, 2018). However, cortical thickness has been linked to the length of CAG repeats in the androgen receptor gene - a measure which influences androgen receptor sensitivity (Raznahan et al., 2010; Zitzmann, 2009). In our analysis, cortical thickness was lower in males in portions of the frontal, temporal and cingulate cortices. On average, the brains of males also expressed a significantly higher gyrification index than brains of females in all but 4 regions examined (the left cuneus cortex, left pericalcarine cortex, and left and right frontal poles) indicating a global, consistent association between sex and gyrification in adolescence. Neuroimaging of adolescents with sex chromosome aneuploidies has revealed both sex differences in gyrification and sex chromosome dosage effects on gyrification, offering insight into a potential genetic explanation for our findings (Fish et al., 2017).

In addition to gray matter macrostructure, the current study examined white matter microstructure.

While previous studies have reported sex differences in FA and MD in the ABCD sample (Lawrence et al., 2021), the current study aimed to further investigate the degree to which gender may have unique or combined effects in addition to the noted sex effects. The sex model (M1) captured the most variance in 79% of white matter regions. Previous research suggests sex differences in white matter microstructure may be related to sex differences in androgen and estrogen. In adolescents, regional FA is correlated with circulating androgen and estradiol (Herting et al., 2012). Additionally, studies of hormone replacement therapy in adults have reported reductions in MD and increases in FA with testosterone administration, while estradiol and antiandrogen therapy resulted in increases in MD and decreases in FA (Kranz et al., 2017). On the other hand, one of the genes involved in myelination, proteolipid protein (PLP1), is located on the X chromosome, is subject to X-inactivation, and has been implicated in multiple sclerosis - a demyelinating disease with a known sex disparity (Cloake et al., 2018). Therefore, more research is needed to elucidate the mechanism through which sex differences in white matter develop in childhood.

These current results also highlight the importance of thoughtful model design in neuroscientific studies of sex and gender. The largest effect sizes for sex were seen for lGI suggesting that sex accounts for a larger proportion of the variance in lGI than it does in any other metric examined. The maximum proportion of variance accounted for by sex was 7.7% in the right superior temporal gyrus. The amount of additional variance captured compared to the null model was also relatively modest, with a median of 0.3% more variance captured by the best model across all ROIs. Furthermore, all of our effect sizes are considered small according to Cohen’s heuristic (e.g., small: *f*^2^≥ 0.02, medium: *f*^2^≥ 0.15, large: *f*^2^ ≥ 0.35; Cohen, 1988), and most are below the average effect size in the ABCD study (0.05; Owens et al., 2021). Therefore, in contrast to the conclusion of Brennan *et al*., our results indicate that sex differences account for a small amount of variance in brain structure, rather than ‘sexual dimorphism’, at ages 9-11 years old. As such, our findings bolster previous assertions that sex differences in the brain do exist, but do not amount to two distinct, non-overlapping sex phenotypes in the brain (Eliot et al., 2021; Friedrichs & Kellmeyer, 2022; Hyde et al., 2018; Rippon et al., 2014). In contrast to sex-related variance, we did not discover any significant neuroanatomical variability related to gender diversity alone. That is, while gender improved model-fit in some regions by accounting for more variance than sex alone, felt-gender was not found to be statistically significant in those models. Similarly, a prior study in adolescents aged 12-17 found no relationship between gender dysphoria and cortical morphometry (Skorska et al., 2021). Though these findings suggest that there is not a direct association between felt-gender or dysphoria and brain structure, the relationship between gender diversity and health outcomes is theorized to be complex and modified by distal and proximal stressors, as well as protective factors (K. K. Tan et al., 2020; Testa et al., 2015). Therefore, further study is needed to explore whether other facets of gender and gender-related aspects of the social environment interact with neurological development. Collectively, our analyses demonstrate that sex - but not felt-gender - accounts for a small fraction of the total structural variance at ages 9-11 years-old.

### 4.1 Limitations

The generalizability of our results is limited by non-response and selection bias. The group of participants excluded due to missing data on the gender questionnaire was more frequently Hispanic, Black, had more advanced pubertal development, and had lower parental education than the included sample. Other authors using ABCD data have also demonstrated that sample sociodemographic composition can be affected by common exclusion criteria, such as image quality, limiting the generalizability of results (Cosgrove et al., 2022). Hence, our exclusion of participants based on image quality, atypical neuroanatomical findings, or missing data may have contributed to bias in our final sample. Finally, because the gender questionnaire was not administered at the baseline imaging visit, we relied on responses collected approximately one year after images were collected.

Therefore, our models assume that felt-gender is relatively stable between the ages of 9-11 years.

Furthermore, this study is a cross-sectional, observational study of sex and gender in humans, and therefore, we cannot ascertain the direction, mechanism, or cause of our results. As such, our findings should not be interpreted as evidence that sex causes structural differences. The concept of permanent, dimorphic sex or gender phenotypes in the brain is at odds with the dynamic nature of sex, gender, and the brain (Friedrichs & Kellmeyer, 2022). Brain structure in general is plastic and can be altered temporarily or permanently by various experiences and environmental factors (Kolb & Gibb, 2011). Social factors like family and school support are also known to mitigate the relationship between gender diversity and mental health problems (Ross-Reed et al., 2019). Our research was limited in scope and not designed to encapsulate all the myriad dimensions of sex and gender that could interact with brain structure. While androgens and estrogens have a documented relationship with brain structure, levels fluctuate with time of day, menstrual cycle phase, and contraceptive use (Uban et al., 2018). Given that hormone collection in ABCD is not consistent with regard to time of day or menstrual cycle phase (Herting et al., 2021; Uban et al., 2018), we chose not to include measures of circulating hormones in our analysis. In terms of genetics, although we were able to detect the presence of a Y chromosome through use of the X and Y allele frequency ratio, we did not examine other sources of sex chromosome variability - including aneuploidy, mosaicism, X-chromosome inactivation, and inactivation escape (Raznahan & Disteche, 2021). Though we failed to reject the null hypothesis of no relationship between felt-gender diversity and brain structure, our results cannot rule out an association between brain structure and other aspects of gender. Moreover, it may be necessary to consider moderating factors such as stress, social exclusion, and victimization (Loso et al., 2023; K. K. Tan et al., 2020). Future work should explore how other gender-related variables - such as gender expression, gender discrimination, and social support - may contribute to brain structure during development. Thus, our analyses should be viewed as an initial step towards disentangling the sex and gender in neurodevelopmental research.

### 4.2 Conclusions

This study demonstrated that variance in white matter microstructure, subcortical volume, cortical thickness and gyrification was related to sex, but not gender diversity in children aged 9-11. Although adding gender diversity increased the amount of outcome variance explained by the model in 8 gray matter ROIs and the addition of the sex-by-gender diversity interaction term improved the model fit in one white matter ROI, neither gender diversity, nor the interaction term were significant in any of these regions. In contrast, the sex model captured the most variance in 76% of the ROIs examined. However, the effect sizes of these sex differences across a variety of macro and microstructural measures were universally small. This suggests that sex accounts for a small proportion of neuroanatomical differences in late childhood.

## Acknowledgments

Data used in the preparation of this article were obtained from the Adolescent Brain Cognitive Development^SM^ (ABCD) Study (https://abcdstudy.org), held in the NIMH Data Archive (NDA). This is a multisite, longitudinal study designed to recruit more than 10,000 children ages 9-11 and follow them over 10 years into early adulthood. The ABCD Study® is supported by the National Institutes of Health and additional federal partners under award numbers U01DA041048, U01DA050989, U01DA051016, U01DA041022, U01DA051018, U01DA051037, U01DA050987, U01DA041174, U01DA041106, U01DA041117, U01DA041028, U01DA041134, U01DA050988, U01DA051039, U01DA041156, U01DA041025, U01DA041120, U01DA051038, U01DA041148, U01DA041093, U01DA041089, U24DA041123, U24DA041147. A full list of supporters is available at https://abcdstudy.org/federal-partners.html. A listing of participating sites and a complete listing of the study investigators can be found at https://abcdstudy.org/consortium_members/. ABCD consortium investigators designed and implemented the study and/or provided data but did not necessarily participate in the analysis or writing of this report. This manuscript reflects the views of the authors and may not reflect the opinions or views of the NIH or ABCD consortium investigators. This study was also funded by RF1MH123223.

## Author Contributions

**Carinna Torgerson:** Conceptualization, Formal analysis, Investigation, Methodology, Writing - original draft. **Hedyeh Ahmadi:** Methodology, Formal analysis, Writing - review & editing. **Jeiran Choupan:** Conceptualization, Funding acquisition, Supervision, Writing - review & editing. **Chun Fan:** Methodology, Writing - review & editing. **John Blosnich:** Methodology, Writing - review & editing. **Megan Herting:** Conceptualization, Funding acquisition, Supervision, Writing - review & editing.

## Data and Code Availability Statement

Data from the ABCD Study Data Release is available for access through the NIMH Data Archive (NDA 3.0 data release 2020: https://dx.doi.org/10.15154/1520591; NDA 4.0 data release 2021; https://dx.doi.org/10.15154/1523041). Only researchers with an approved NDA Data Use Certification (DUC) can obtain ABCD Study data.

