## Supplemental Methods for "Sex, gender diversity, and brain structure in children ages 9 to 11 years old"

### **Figure 1.**

*A graphical representation of exclusionary criteria and the quality control process for the study.*

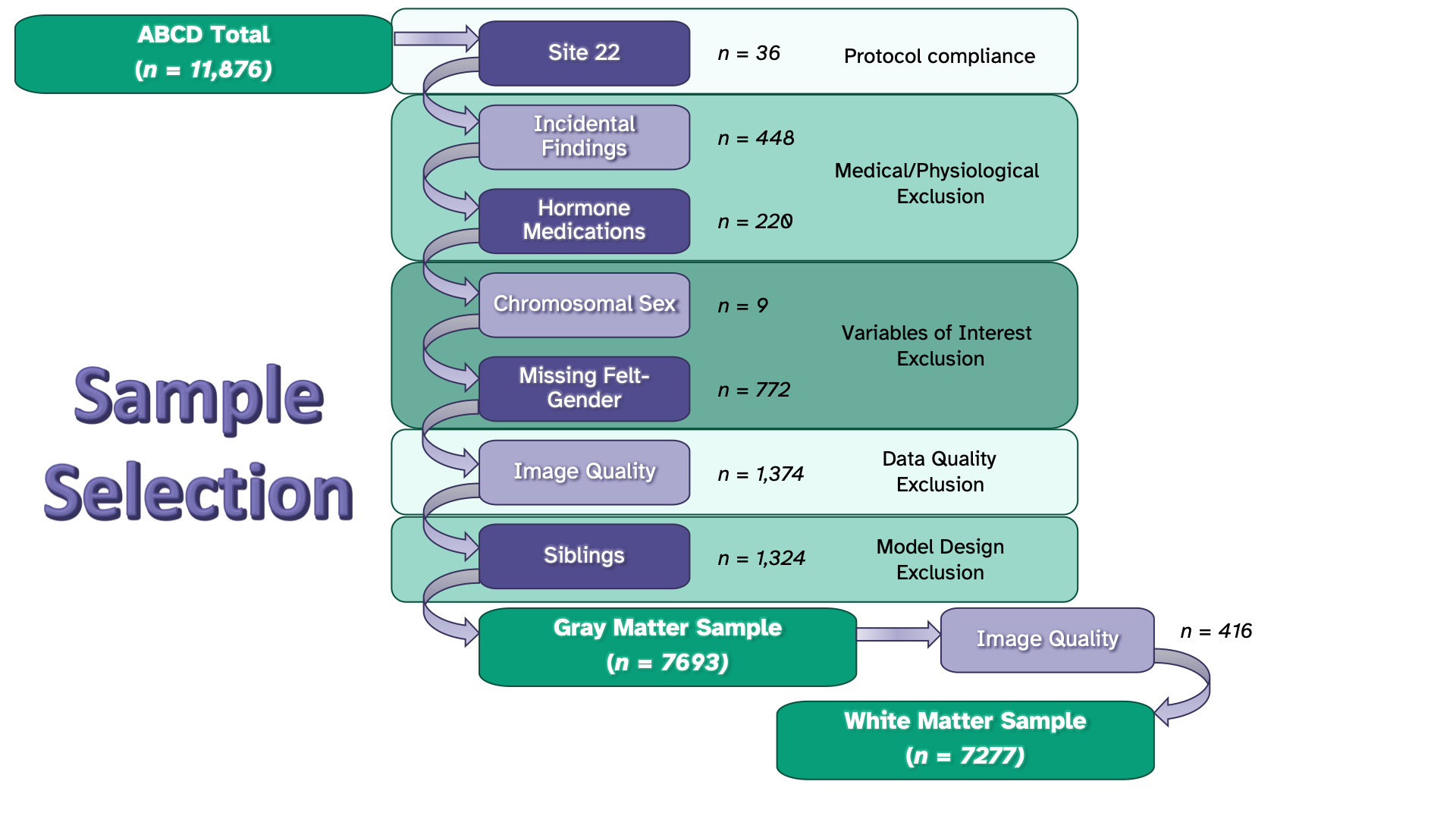

### **Table 1:**

*List of hormone medications excluded from the sample.*

| **RxNormNum** | **ATC L4 Chemical Subgroup** | **ABCD_MedName** |
| --- | --- | --- |
| 3251 | H01BA - Vasopressin and analogues | Desmopressin |
| 5492 | H02AB - Glucocorticoids | Hydrocortisone |
| 8640 | H02AB - Glucocorticoids | Prednisone |
| 10582 | H03AA - Thyroid hormones | Thyroxine |
| 10759 | H02AB - Glucocorticoids | Triamcinolone |
| 10761 | H02AB - Glucocorticoids | Triamcinolone Acetonide |
| 27199 | H02AB - Glucocorticoids | Hydrocortisone valerate |
| 42634 | H01BA - Vasopressin and analogues | Desmopressin Acetate |
| 103403 | H02AB - Glucocorticoids | Hydrocortisone 10 MG/ML Topical Lotion |
| 134920 | H02AB - Glucocorticoids | Maxitrol |
| 197123 | H01AC - Somatropin and somatropin agonists | Humatrope |
| 202829 | H01AC - Somatropin and somatropin agonists | Norditropin |
| 203658 | H01BA - Vasopressin and analogues | Vasopressin |
| 206511 | H02AB - Glucocorticoids | Hydrocortisone 10 MG/ML / Neomycin 3.5 MG/ML / Polymyxin B 10000 UNT/ML Otic Solution [Cortisporin] |
| 218002 | H03AA - Thyroid hormones | Levoxyl |
| 224920 | H03AA - Thyroid hormones | Synthroid |
| 316582 | H02AB - Glucocorticoids | Prednisone 20 MG |
| 371716 | H01BA - Vasopressin and analogues | Desmopressin Oral Tablet |
| 372595 | H03AA - Thyroid hormones | Levothyroxine Oral Tablet |
| 374371 | H01BA - Vasopressin and analogues | Desmopressin Nasal Spray |
| 377712 | H02AB - Glucocorticoids | Hydrocortisone Topical Cream |
| 377831 | H02AB - Glucocorticoids | Triamcinolone Topical Cream |
| 404930 | H02AB - Glucocorticoids | Ciprodex |
| 567806 | H03BB - Sulfur-containing imidazole derivatives | Methimazole 5 MG [Tapazole] |
| 644964 | H01AC - Somatropin and somatropin agonists | Omnitrope |
| 833007 | H01BA - Vasopressin and analogues | Desmopressin Acetate 0.2 MG |
| 847245 | H01AC - Somatropin and somatropin agonists | 1.5 ML Somatropin 10 MG/ML Pen Injector [Norditropin] |
| 849517 | H01BA - Vasopressin and analogues | Desmopressin Acetate 0.1 MG Oral Tablet [DDAVP] |
| 849524 | H01BA - Vasopressin and analogues | Desmopressin Acetate 0.2 MG Oral Tablet [DDAVP] |
| 903316 | H03AA - Thyroid hormones | Levothyroxine Sodium 0.025 MG / liothyronine sodium 0.00625 MG Oral Tablet [Thyrolar] |
| 966249 | H03AA - Thyroid hormones | Levothyroxine Sodium 0.175 MG Oral Tablet |
| 966394 | H03AA - Thyroid hormones | Levothyroxine Sodium 0.1 MG [Synthroid] |
| 1007849 | H02AB - Glucocorticoids | Triamcinolone / Urea |
| 1085636 | H02AB - Glucocorticoids | Triamcinolone Acetonide 0.001 MG/MG Topical Ointment |
| 1110806 | H02AB - Glucocorticoids | Triamcinolone Acetonide 0.0005 MG/MG Topical Ointment [Trianex] |
| 1164010 | H02AB - Glucocorticoids | Hydrocortisone Pill |
| 1170903 | H02AB - Glucocorticoids | Ciprodex Otic Product |
| 1185122 | H01AC - Somatropin and somatropin agonists | Norditropin Injectable Product |
| 1186074 | H03AA - Thyroid hormones | Synthroid Pill |
| 1250490 | H02AB - Glucocorticoids | Hydrocortisone acetate 5 MG/ML Topical Cream |
| 1648996 | H02AB - Glucocorticoids | Triamcinolone Acetonide 0.25 MG/ML [Triderm] |

### **Table 2.**

*Questions from the felt-gender dimension ABCD youth gender questionnaire completed by participants.*

| **Question** | **XX version** | **XX Scoring** | **XY version** | **XY scoring** |
| --- | --- | --- | --- | --- |
| **1** | How much do you feel like a **girl**? | 5 = Totally  4 = Mostly  3 = Somewhat  2 = A little  1 = Not at all | How much do you feel like a **boy**? | 5 = Totally  4 = Mostly  3 = Somewhat  2 = A little  1 = Not at all |
| **2** | How much do you feel like a **boy**? | 1 = Totally  2 = Mostly  3 = Somewhat  4 = A little  5 = Not at all | How much do you feel like a **girl**? | 1 = Totally  2 = Mostly  3 = Somewhat  4 = A little  5 = Not at all |

### **Table 3.**

*A complete list of ROIs to be assessed. (L/R) denotes regions with separate ROIs for the left and right hemisphere.*

| ***FA and MD*** | ***Volume*** | ***CTh and lGI*** | |
| --- | --- | --- | --- |
| ***Limbic*** | ***Basal Ganglia*** | ***Frontal*** | ***Parietal*** |
| *Fornix (L/R)* | *Caudate (L/R)* | *Superior Frontal (L/R)* | *Superior Parietal (L/R)* |
| *Cingulate Cingulum (L/R)* | *Putamen (L/R)* | *Caudal Middle Frontal (L/R)* | *Inferior Parietal (L/R)* |
| *Parahippocampal Cingulum (L/R)* | *Pallidum (L/R)* | *Rostral Middle Frontal (L/R)* | *Supramarginal (L/R)* |
| ***Projection*** | *Accumbens (L/R)* | *Pars Opercularis (L/R)* | *Postcentral (L/R)* |
| *Corticospinal/Pyramidal Tracts (L/R)* | ***Limbic*** | *Pars Orbitalis (L/R)* | *Precuneus (L/R)* |
| *Anterior Thalamic Radiations (L/R)* | *Amygdala (L/R)* | *Pars Triangularis (L/R)* | ***Occipital*** |
| ***Association*** | *Hippocampus (L/R)* | *Lateral Orbitofrontal (L/R)* | *Lateral Occipital (L/R)* |
| *Uncinate Fasciculus (L/R)* | *Thalamus (L/R)* | *Medial Orbitofrontal (L/R)* | *Lingual (L/R)* |
| *Superior Longitudinal Fasciculus (L/R)* | ***Corpus Callosum*** | *Precentral Gyrus (L/R)* | *Cuneus (L/R)* |
| *Inferior Longitudinal Fasciculus (L/R)* | *Anterior CC* | *Paracentral Gyrus (L/R)* | *Pericalcarine (L/R)* |
| *Inferior Fronto-Occipital Fasciculus (L/R)* | *Mid Anterior CC* | *Frontal Pole (L/R)* | ***Cingulate*** |
| ***Callosal*** | *Central CC* | ***Temporal*** | *Rostral Anterior Cingulate (L/R)* |
| *Forceps Major* | *Mid Posterior CC* | *Superior Temporal (L/R)* | *Caudal Anterior Cingulate (L/R)* |
| *Forceps Minor* | *Posterior CC* | *Middle Temporal (L/R)* | *Posterior Cingulate (L/R)* |
| *Corpus Callosum* | ***Other*** | *Inferior Temporal (L/R)* | *Isthmus Cingulate (L/R)* |
|  | *Cerebellum Cortex (L/R)* | *Banks of the Superior Temporal Sulcus (L/R)* | ***Insula*** |
|  | *Brain Stem* | *Fusiform (L/R)* | *Insula (L/R)* |
|  |  | *Transverse Temporal (L/R)* |  |
|  |  | *Entorhinal (L/R)* |  |
|  |  | *Temporal Pole (L/R)* |  |
|  |  | *Parahippocampal (L/R)* |  |

### **Table 4.**

The linear mixed effects model equations for the null model (M0) and the four sex and gender models.

| **M0** | ROI*_i_* ∼ *N* (*μ*, *σ*^2^)  *μ* = *α_j_*_[_*_i_*_]_ + *β*_1_(PDS_Early_) + *β*_2_(PDS_Mid/Late_) + *β*_3_(Age) + *β*_4_(Edu_HS or GED_) + *β*_5_(Edu_Some College_) + *β*_6_(Edu_Bachelor_) + *β*_7_(Edu_Post Graduate Degree_) + *β*_8_(Race_Black_) + *β*_9_(Race_Multiracial Black_) + *β*_10_(Race_Multiracial Non-Black_) + *β*_11_(Race_Other_) + *β*_12_(Ethnicity_Hispanic_) + *β*_13_(Scanner_GE_) + *β*_14_(Scanner_Philips_)  *α_j_* ∼ *N* (*μ_αj_* ,*σ*^2^*_αj_*), for Site j = 1,…,21  Reference categories: Sex_XX_ ; Puberty_Pre_ ; Edu_< HS Diploma_ ; Race_White_ ; Ethnicity_Non-Hispanic_ ; Scanner_Siemens_ |
| --- | --- |
| **M1** | ROI*_i_* ∼ *N* (*μ*, *σ*^2^)  *μ* = *α_j_*_[_*_i_*_]_ + ***β*_1_(Sex_XY_)** + *β*_2_(PDS_Early_) + *β*_3_(PDS_Mid/Late_) + *β*_4_(Age) + *β*_5_(Edu_HS or GED_) + *β*_6_(Edu_Some College_) + *β*_7_(Edu_Bachelor_) + *β*_8_(Edu_Post Graduate Degree_) + *β*_9_(Race_Black_) + *β*_10_(Race_Multiracial Black_) + *β*_11_(Race_Multiracial Non-Black_) + *β*_12_(Race_Other_) + *β*_13_(Ethnicity_Hispanic_) + *β*_14_(Scanner_GE_) + *β*_15_(Scanner_Philips_)  *α_j_* ∼ *N* (*μ_αj_* ,*σ*^2^*_αj_*), for Site j = 1,…,21  Reference categories: Sex_XX_ ; Puberty_Pre_ ; Edu_< HS Diploma_ ; Race_White_ ; Ethnicity_Non-Hispanic_ ; Scanner_Siemens_ |
| **M2** | ROI*_i_* ∼ *N* (*μ*, *σ*^2^)  *μ* = *α_j_*_[_*_i_*_]_ + ***β*_1_(Felt-gender)** + *β*_2_(PDS_Early_) + *β*_3_(PDS_Mid/Late_) + *β*_4_(Age) + *β*_5_(Edu_HS or GED_) + *β*_6_(Edu_Some College_) + *β*_7_(Edu_Bachelor_) + *β*_8_(Edu_Post Graduate Degree_) + *β*_9_(Race_Black_) + *β*_10_(Race_Multiracial Black_) + *β*_11_(Race_Multiracial Non-Black_) + *β*_12_(Race_Other_) + *β*_13_(Ethnicity_Hispanic_) + *β*_14_(Scanner_GE_) + *β*_15_(Scanner_Philips_)  *α_j_* ∼ *N* (*μ_αj_* ,*σ*^2^*_αj_*), for Site j = 1,…,21  Reference categories: Sex_XX_ ; Puberty_Pre_ ; Edu_< HS Diploma_ ; Race_White_ ; Ethnicity_Non-Hispanic_ ; Scanner_Siemens_ |
| **M3** | ROI*_i_* ∼ *N* (*μ*, *σ*^2^)  *μ* = *α_j_*_[_*_i_*_]_ + ***β*_1_(Sex_XY_) + *β*_2_(Felt-gender)** + *β*_3_(PDS_Early_) + *β*_4_(PDS_Mid/Late_) + *β*_5_(Age) + *β*_6_(Edu_HS or GED_) + *β*_7_(Edu_Some College_) + *β*_8_(Edu_Bachelor_) + *β*_9_(Edu_Post Graduate Degree_) + *β*_10_(Race_Black_) + *β*_11_(Race_Multiracial Black_) + *β*_12_(Race_Multiracial Non-Black_) + *β*_13_(Race_Other_) + *β*_14_(Ethnicity_Hispanic_) + *β*_15_(Scanner_GE_) + *β*_16_(Scanner_Philips_)  *α_j_* ∼ *N* (*μ_αj_* ,*σ*^2^*_αj_*), for Site j = 1,…,21  Reference categories: Sex_XX_ ; Puberty_Pre_ ; Edu_< HS Diploma_ ; Race_White_ ; Ethnicity_Non-Hispanic_ ; Scanner_Siemens_ |
| **M4** | ROI*_i_* ∼ *N* (*μ*, *σ*^2^)  *μ* = *α_j_*_[_*_i_*_]_ + ***β*_1_(Sex_XY_) + *β*_2_(Felt-gender)** + *β*_3_(PDS_Early_) + *β*_4_(PDS_Mid/Late_) + *β*_5_(Age) + *β*_6_(Edu_HS or GED_) + *β*_7_(Edu_Some College_) + *β*_8_(Edu_Bachelor_) + *β*_9_(Edu_Post Graduate Degree_) + *β*_10_(Race_Black_) + *β*_11_(Race_Multiracial Black_) + *β*_12_(Race_Multiracial Non-Black_) + *β*_13_(Race_Other_) + *β*_14_(Ethnicity_Hispanic_) + *β*_15_(Scanner_GE_) + *β*_16_(Scanner_Philips_) + ***β*_17_(Felt-gender * Sex_XY_)**  *α_j_* ∼ *N* (*μ_αj_* ,*σ*^2^*_αj_*), for Site j = 1,…,21  Reference categories: Sex_XX_ ; Puberty_Pre_ ; Edu_< HS Diploma_ ; Race_White_ ; Ethnicity_Non-Hispanic_ ; Scanner_Siemens_ |

### **Figure 2.**

*A graphical representation of criteria used to determine the best-fitting model for each region. The (>) symbol indicates that an ANOVA comparison of the two models yielded a p < 0.05. M0 = null model (i.e., covariates only); M1 = null model + sex; M2 =null model + gender; M3 = null model + sex + gender; M4 = null model + sex + gender + sex-by-gender interaction; AIC = Akaike Information Criterion*

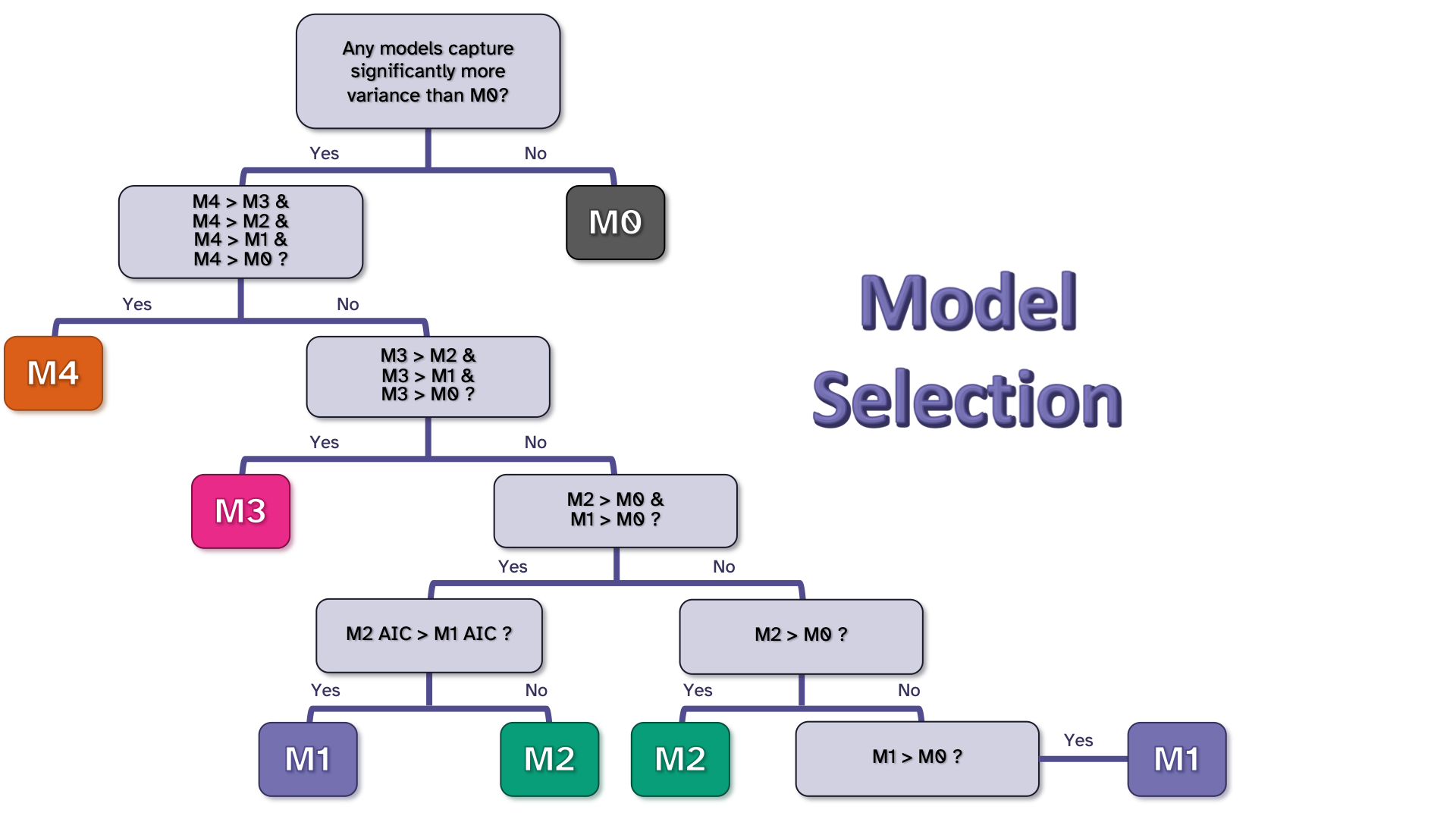
